# Initiation site of experimentally-evoked spreading depolarizations influence tissue outcomes in a murine stroke model

**DOI:** 10.1101/2025.10.03.680372

**Authors:** Michael C. Bennett, Russell A. Morton, Andrew P. Carlson, C. William Shuttleworth

## Abstract

Spreading depolarization waves (SDs) are implicated in secondary expansion of brain injuries and are the target for initial clinical intervention trials. However, the assumption that SD directly causes neuronal injury has been challenged by recent findings with experimentally-induced SD in stroke models. The current study addressed this controversy by examining whether stroke consequences are confounded by the precise location of experimental SD initiation. Focal ischemic lesions were generated by transient distal middle cerebral artery occlusion in male mice. Clusters of SDs (6 at 10-min intervals) were induced by either focal KCl application or optogenetic stimulation during occlusion. SDs were initiated either in regions close to the infarct core (penumbral-SD; <50% perfusion) or in less compromised tissue in the same hemisphere (remote-SD; >70% perfusion). Despite the fact that all SDs fully invaded stroke expansion areas, the location of experimental SD induction had significant effects on stroke outcomes measured 48 hours after reperfusion. Penumbral-SDs resulted in larger infarct expansion than seen in control stroke mice lacking experimentally-imposed SD. Conversely, remote-SDs led to significantly smaller infarcts than stroke alone. Laser speckle contrast imaging of blood flow in injury expansion areas showed enhanced hypoperfusion responses after penumbral-SDs and larger hyperemic responses after remote-SDs, suggesting that differential vascular responses could contribute to stroke outcomes. Overall, this study helps to reconcile different prior reports by showing that experimentally-induced SDs can either exacerbate or reduce stroke-induced injury depending on the SD initiation site and further strengthens evidence for injurious roles of SDs initiating in vulnerable brain tissue.

## Introduction

Spreading depolarizations (SD) are slowly propagating waves of near-complete neuronal and glial depolarization that lead to transient suppression of neuronal activity and large vascular responses (Leao, 1944; Somjen, 2001). SDs result in a high metabolic burden and can have detrimental consequences when they occur in tissue lacking metabolic supply, such as in ischemic stroke (von Bornstädt et al., 2015; Hartings et al., 2017). SDs are recorded in peri-infarct areas for multiple days after an ischemic insult, and are strongly implicated in secondary injury and lesion expansion (Busch et al., 1996; Hartings et al., 2003; Dohmen et al., 2008; Nakamura et al., 2010). Preclinical characterization of SD has identified neuronal excitotoxicity, edema, and inversed neurovascular coupling as mechanisms that can contribute to injury (Aiba and Shuttleworth, 2012; Hinzman et al., 2015; von Bornstädt et al., 2015; Reinhart et al., 2023) and preclinical work has informed recent clinical interventions and trials examining whether targeting SDs in the clinic is beneficial to patient outcomes (Carlson et al., 2023; Hartings et al., 2023). Part of the evidence supporting a role of SD in ischemic infarct expansion comes from studies where an additional burden of experimentally-evoked SDs has been applied in a rodent stroke model. For example, step-wise increases in infarct size accompanied SDs initiated with focal KCl (Busch et al., 1996). However, a recent study using optogenetic SD induction in a stroke model showed no additional infarct expansion, questioning both prior work with KCl stimulation and more generally the causal role of SD in stroke expansion (Sugimoto et al., 2023, 2024). This discrepancy is important to address, especially as SD-targeting is moving into clinical trials.

The consequences of SD occur along a continuum from responses in healthy, metabolically supplied tissue (where brain tissue can fully recover quite quickly), to ischemic areas where the extended depolarization and disrupted neurovascular coupling can lead to irrecoverable neuronal injury (Pietrobon and Moskowitz, 2014; Dreier and Reiffurth, 2015; Hartings et al., 2017). It has been shown that differences in the spread of spontaneously-occurring SDs impact the expansion of ischemic infarcts (Binder et al., 2022). When SDs and the associated spreading hyperemia extended from the metabolically vulnerable stroke penumbra, infarct size was increased compared to SDs that propagated across the entire hemisphere. The ischemic penumbra is a “hot-zone” for SD initiation in stroke, where SD can be triggered due to metabolic supply/demand mismatch (e.g. transient hypotension or somatosensory activation (von Bornstädt et al., 2015; Carlson et al., 2023). While it is thus likely that SDs that occur in stroke progression initiate at these vulnerable locations, experimental studies that induce an additional burden of SDs evoked experimentally with focal KCl or optogenetic stimulation may not initiate the events in the same location. For practical experimental reasons, SDs are often initiated in more remote locations and allowed to propagate into the vulnerable penumbra (e.g. Reinhart et al., 2023) or intermediate locations.

In the present study, we sought to examine whether the damaging effects of experimentally induced SD in the ischemic brain are highly dependent on the site of SD initiation. We utilized a focal, transient model of ischemic stroke, closely following the methods of recent studies (Sugimoto et al., 2023, 2024), to test the hypothesis that SDs experimentally induced within metabolically depleted, peri-infarct “hot zones” would cause lesion growth, but SDs initiated outside this critical area would not cause secondary injury. We found that, regardless of the SD initiating stimulus used (KCl or optogenetic), SDs initiated in penumbral regions caused infarct expansion. In contrast, SDs initiated in remote regions invaded the same infarct expansion areas, but exhibited an increase in cerebral perfusion and significantly smaller infarcts compared to stroke brain naïve to SD generation. Our results help reconcile the recent debate on whether SDs contribute to ischemic stroke injury and emphasize the importance of refinement of experimental approaches by taking the initiation site and vascular dynamics into account.

## Materials and Methods

### Animals

All animal procedures were performed in accordance with protocols approved by the UNM Health Sciences Center Institutional Animal Care and Use Committee. Adult male mice (10 – 15-weeks old; 4/cage) were used for all experiments and were housed in standard, ventilated cages with *ad libitum* food and water access on a 12-hour light/dark cycle. C57Bl/6 mice were used for KCl stimulation of SD. For optogenetic SD initiation, transgenic mice expressing channelrhodopsin-2 under the Thy1 promoter were used from in-house offspring bred from a transgenic line purchased from The Jackson Laboratory (B6-Tg(Thy1-COP4[EYFP])/J; Jax #007612) (Pinkowski et al., 2024).

### Surgical preparation and transient distal middle cerebral artery occlusion

Isoflurane (4% induction and 1.5% maintenance) was used for anesthesia in all experiments as described previously (Sugimoto et al., 2023). Briefly, body temperature was maintained between 36.5 – 37⁰C by using a feedback-controlled heating pad (Kent Scientific Corporation). Mice were then placed in a stereotaxic frame with supplemental oxygen (70% N_2_ and 30% O_2_) provided via a nose cone and sterile eye ointment was applied (OptixCare, Country Side Pet). To facilitate vascular imaging, the skull was exposed by a midline incision and kept moist with saline (0.9% NaCl) or mineral oil to improve image quality.

For a focal stroke, an incision was made in the skin between the orbit and external auditory cortex before the temporalis muscle was resected. A small craniotomy (∼2mm) was then drilled in the temporal bone immediately rostral to the zygomatic arch and squamosal bone to reveal the distal middle cerebral artery (MCA). MCA occlusion (MCAo) was performed just proximal to the branching of the MCA via a microvascular clip as previously described (Figure 1B) (Buchan et al., 1992; Sugimoto et al., 2023). Laser speckle contrast imaging (LCSI, see below) was used to map the extent of perfusion deficit and confirm successful reperfusion after 60 min occlusion. All mice in the study showed similar degree of perfusion deficit (Figure 1C). Experimental SDs were initiated during the occlusion period (see below). After unclipping and confirmation of successful reperfusion, the skin was then sutured and following surgical procedure mice were given analgesic (buprenorphine; 0.25ml/25g) immediately after surgery and then every 12 hours until they were sacrificed 48 hours post-occlusion.

**Figure 1:**
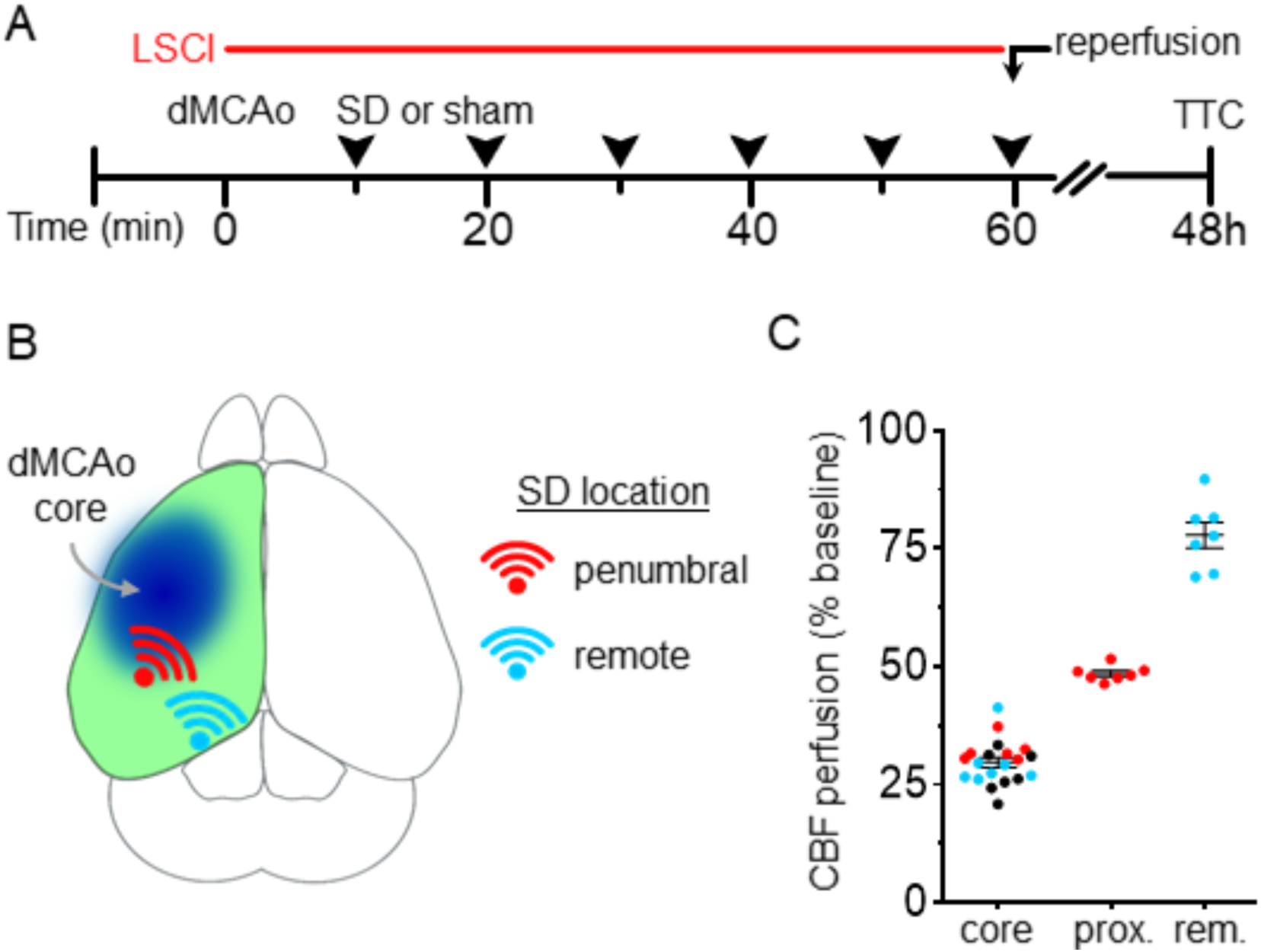
Experimental SD initiation sites, at different locations relative to area of perfusion deficit from dMCAo. (A) Timeline of experiments. Prior to transient distal MCA occlusion (dMCAo), LSCI imaging was performed for baseline comparisons and then continued immediately after dMCAo for the remainder of the experiments. Burr holes were also prepared for SD induction (arrowheads; 10 minutes between each SD). At 60 minutes, occlusion was reversed and reperfusion was confirmed. Mice were sacrificed 48 hours post-MCAo and brains were collected for infarct evaluation with TTC staining. (B) Diagram showing the approximate locations of experimental SD initiation, at sites either penumbral (red) or remote (blue) to the ischemic core. SDs were experimentally evoked by focal KCl (KCl-SD) (C) Group data (normalized to pre-MCAo baselines) showing perfusion deficits within the core infarct area, and at the two different initiation sites.

### Laser speckle contrast imaging

Laser speckle contrast imaging (LSCI) of relative blood flow began with a 3-minute baseline recording prior to MCA occlusion. Immediately following MCAo, mice were transferred to LSCI setup and monitored continuously for the next 60 minutes to observe CBF changes before, during and after SDs. The exposed skull was illuminated with a 785nm laser diode (Thorlabs) and backscattered light was then passed through a long pass filter (720nm) and focused with an SLR camera lens (55mm, f-stop = 5.6, Nikon). LSCI images were collected at ∼4 Hz via a digital CCD camera (Stingray F-504B; Allied Vision Technologies). The perfusion maps were then calculated and displayed in real time utilizing LabVIEW software that had been previously modified from Bernard Choi, UCI (Yang et al., 2011). CBF in regions of interest (ROIs) was assessed offline in Fiji (RRID:SCR_002285) and compared with matched ROIs in the ipsilateral hemisphere.

### Experimental SD initiation

Experimentally-induced SDs were generated as a cluster of 6 events at 10 min intervals, beginning following occlusion of the MCA (after 10 min). Occlusion of the MCA usually causes spontaneous initiation of an SD (Shin et al., 2006; Reinhart et al., 2023). Due to technical limitations (30 – 45 seconds for repositioning of the animal for imaging) the onset of these initial events was not detected, but LSCI captured their termination in most animals (82.1%; n=32/39). Spontaneous SDs were also recorded immediately following removal of the microvascular clip during reperfusion in a number of mice (56.4%; n=22/39). Other spontaneously-occurring SDs were not observed during the period of experimental SD initiation. All spontaneously SDs propagated outwards from the margin of the ischemic core and occurred across experimental groups.

For KCl-SD induction, burr holes (1mm diameter) were drilled prior to stroke induction in locations proximal and distal to the infarct core. Intermittent cool saline was applied to the skull surface during drilling to minimize heating and LSCI was used to monitor for SD generation during burr hole preparation and was used as criteria for removal from the study. Cortical areas with >50% CBF reduction were deemed “proximal” to infarct core, while areas with <30% reduction (70% perfusion) were deemed “distal.” SDs were initiated by application of 1M KC l(∼1µL) which is known to produce reliable SDs (Reinhart et al., 2023). SDs propagated through the areas of perfusion deficit and were monitored by perfusion changes detected using LSCI (Chung et al., 2018). Excess KCl was removed via saline-soaked cotton swab 60 sec after initial application to reduce likelihood of secondary SD initiation. SDs were initiated every 10 minutes over the next 60 minutes before reperfusion (Figure 1A).

SD was initiated optogenetically via focal illumination through the intact skull with a fiber-coupled LED (470 nm, #M470F4, ThorLabs; 30 seconds at 3mW) connected to an optogenetic patch cable (400 µm core; 0.39NA fiber, ThorLabs). As for KCl-induced SDs, the optical fiber was placed at locations proximal to the perfusion deficit (<50% CBF reduction) or in more remote regions maintaining at least 70% CBF.

### TTC staining

Mice were sacrificed 48-hours post-distal MCAo and brains extracted for infarct measurements. with histochemical analysis using 2,3,5-triphenyltetrazolium chloride (TTC) (Bederson et al., 1986). Briefly, brains were immediately transferred to cold phosphate buffer saline (PBS) before being prepared into 2mm thick coronal slices. To visualize infarcts, slices were then soaked in 2% TTC (Sigma), dissolved in PBS, for 20 – 30 minutes at room temperature. Infarct sizes were measured by calculations done in Fiji (RRID:SCR_002285) software and edema corrected utilizing calculations previously described (Nouraee et al., 2019).

### Experimental Design and Statistical Analysis

Data are reported as mean ± standard deviation. Adult male mice were used for all studies (39 total). Sample sizes for experiments reported below were determined from a set of pilot data and power analyses conducted in G*Power (version 3.1.9.7; Los Angeles, CA) and statistical analyses were performed in GraphPad Prism (version 10.3.9; La Jolla, CA). Parametric tests evaluating statistical significance included unpaired t-tests and one- and two-way analysis of variance (ANOVA). Normal distribution was evaluated with Shapiro-Wilk’s test and statistical significance was identified as P values <0.05, with Bonferroni’s correction used for multiple comparisons.

## Results

### Selection of SD initiation sites

Initial characterization focused on SD initiated with local application of KCl (see Methods). SDs were experimentally initiated every 10 minutes over a 1-hour period of distal MCA occlusion (dMCAo; see Methods). Experimentally-induced SDs were initiated at one of two locations, determined by the degree of perfusion deficit (Figure 1B). SDs experimentally initiated close to the perfusion deficit core (at sites with CBF reduction >50%) are hereafter referred to as “penumbral-SD”, whereas SDs experimentally initiated in less compromised tissue (sites with CBR reduction <30%) are referred to as “remote-SDs”. As noted above (Methods), spontaneously-occurring SDs were observed during occlusion and often during reperfusion after the termination of dMCAO. However, in these studies the main burden of SDs was from experimentally-induced events, as KCl stimulation reliably generated SD with each stimulus, regardless of site of induction (average number of SDs generated for dMCAo alone: 1.4 ± 0.5; remote SDs: 7.2 ± 1.2; penumbral SDs: 7.7 ± 0.95; one-way analysis of variance (ANOVA) p<0.0001; n = 7 mice/group).

### Differences in SD characteristics depending on region of initiation

SDs cause significant hemodynamic changes which follow the depolarization wave (Brennan et al., 2007; Ayata and Lauritzen, 2015) and can be visualized with LSCI (Figure 2). Previous studies have utilized LSCI imaging to assess hemodynamic responses during SD initiation and propagation (Santos et al., 2016; Pacheco et al., 2019). The directionality of SD propagation differed with initiation site. Penumbral-SDs tended to propagate first towards the sagittal sinus and then across the entire hemisphere (Supplemental Movie S1) while remote-SDs propagated rostrally as single wave across the hemisphere (Supplemental Movie S2). However, regardless of initiation site, SDs propagated across the entire hemisphere including infarct expansion zones (Figure 2B).

**Figure 2:**
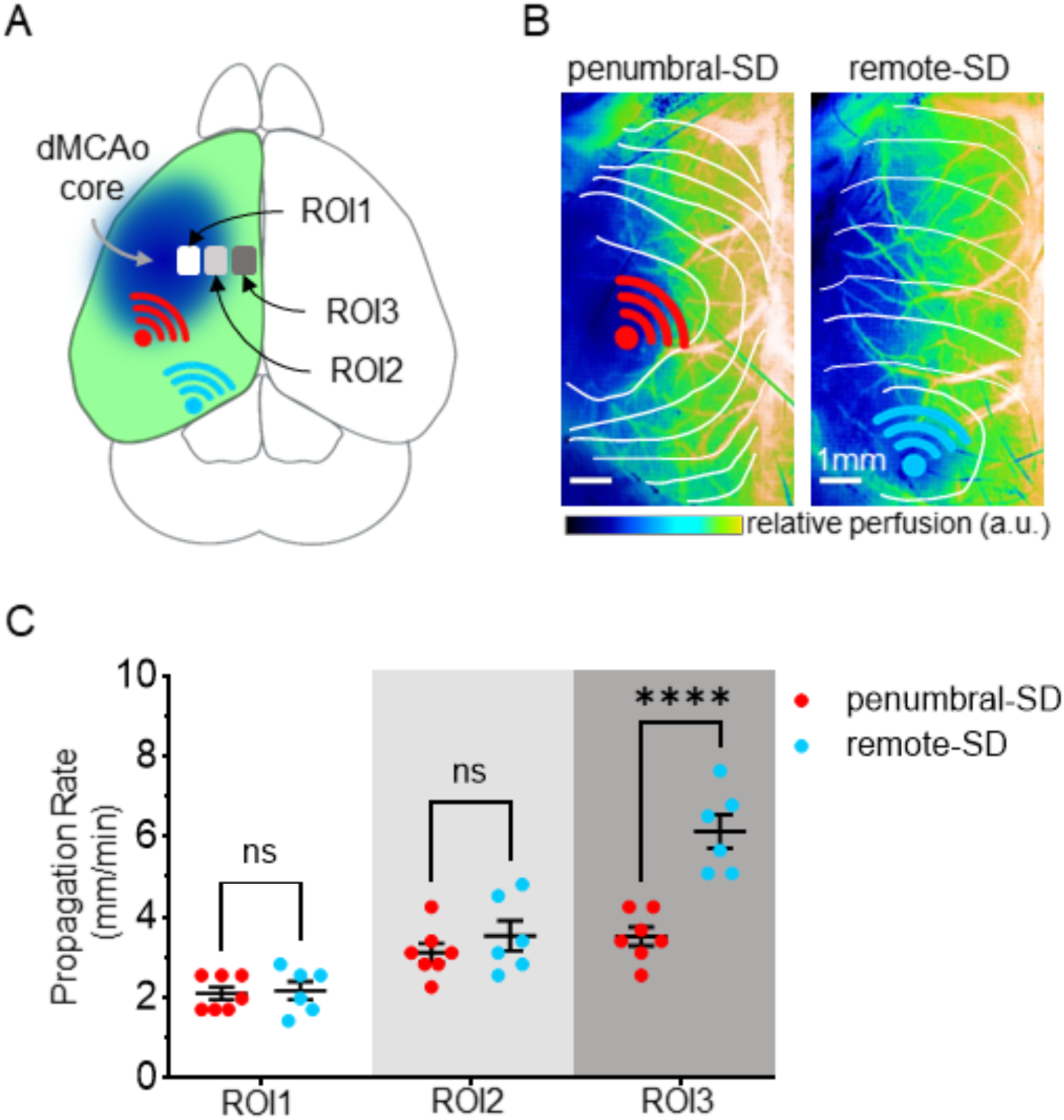
Remote and penumbral SDs invade entire hemisphere. (A) Diagram showing the approximate locations for propagation measurement areas where the perfusion deficit increases from ROI1 (most deficit) to ROI3 (least deficit). (B) Representative pseudo-colored LSCI images showing propagation of experimentally-induced SDs in relation to the perfusion deficit produced by dMCAo. SD initiation sites are indicated by red and blue symbols, and the propagating wavefront illustrated by white lines (15 seconds between each), showing propagation across the entire ipsilateral hemisphere regardless of location of SD initiation (left: penumbral; right: remote) scale bar: 1mm. (C) Summary data of propagation rates at different recording sites illustrated in A (see Methods for definitions of recording sites). Rates were slowest in most ischemic areas (ROI1), as compared to distal regions (ROI3). Remote SDs (blue dots) propagated faster than penumbral SDs, but only in the distal zone (n = 6). Normal distribution was evaluated by Shapiro-Wilks test (B: p = 0.3151). **P < 0.008, ***P < 0.0003, ****P < 0.0001

The propagation rate of SD was also different, based on the site of experimental initiation (Figure 2B). Previous studies emphasize that that the cumulative effects of SDs occurring together in a cluster can lead to damage (Hartings et al., 2003; Dohmen et al., 2008). Therefore, our analyses focused on the last SD in the cluster of 6 SDs delivered experimentally. SDs were allowed to propagate outside of initiation sites (for 10 seconds) to minimize interference of the SD induction method (KCl or optogenetic) on characteristics of SD propagation. Propagation rate was also assessed in infarct expansion zones, identified from pilot studies (comparing perfusion deficits with infarct volumes at 48 hours) as regions with ∼60% baseline CBF, medial to the center of the perfusion deficit. All rates were within the well-known range (2 – 6 mm min^-1^) of SD propagation (Somjen, 2001; Woitzik et al., 2013) but propagation was faster in tissues with greater metabolic capacity, independent of site of SD-induction (in mm min^-1^; peri-infarct zone: 2.1 ± 0.47; medial zone: 3.3 ± 0.77; and distal zone: 4.7 ± 1.6; repeated measures one-way ANOVA, F (12, 24) = 1.962, p<0.0773). Furthermore, SDs initiated in remote regions had faster propagation rates in less compromised distal regions, than SDs initiated in proximal regions that traveled through the same locations. Two-way ANOVA of this grouped data revealed a significant effect of initiation site on propagation rate of remote-SDs in distal, healthy regions compared to penumbral-SDs (6.1 ± 1.02 versus 3.5 ± 0.6 mm min^-1^, respectively; dF = 33, F = 20.53, p<0.0001; Figure 2C).

### SD initiation site modulates stroke expansion 48 hours after reperfusion

Figure 3 shows significant differences in infarct expansion, dependent on the site of experimental SD initiation. Figure 3A shows representative TTC images and Figure 3B shows group data. An additional burden of experimentally-initiated SDs from the penumbral location led to larger infarcts than those in the MCAo alone control group (33.7 ± 3.1 versus 27.5 ± 5.2 mm^3^, respectively; n = 7 mice/group; p = 0.0265) and in the remote-SD group (33.7 ± 3.1 versus 20.8 ± 3.4 mm^3^, respectively; n = 7 mice/group; p<0.0001). Interestingly, remote-SDs led to significantly smaller infarcts than the MCAo alone (20.8 ± 3.4 versus 27.5 ± 5.2 mm^3^, respectively; n = 7 mice/group; p=0.017).

**Figure 3:**
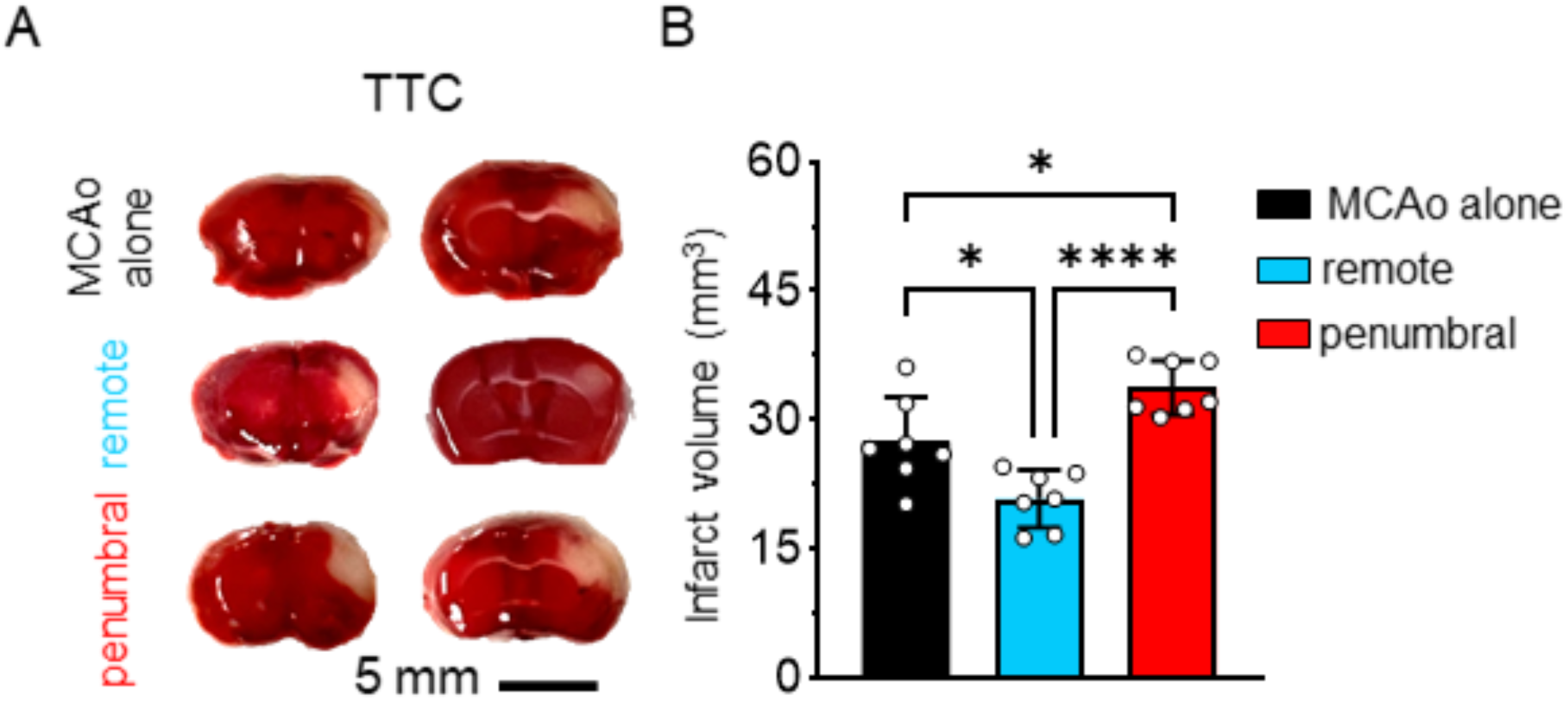
Location of experimental SD initiation affects infarct size. (A) Representative images of brain sections (2mm) stained with 2,3,5-triphenyl tetrazolium chloride (TTC) 48 hours post-MCAo. Representative images from the MCAo alone group (top row), remote SD group (middle) and penumbral SD group (bottom row). (B) Summary data showing that remote SD (blue bar) group had decreased infarct size as compared with control MCAo alone (black bar) and penumbral SD (red bar) groups. Penumbral SD group also had increased infract size as compared with MCAo alone controls (n = 7/group). Normal distribution was evaluated by Shapiro-Wilk test (B: p = 0.7431). *P < 0.03, ****P < 0.0001

One replicate of each of the three experimental SD groups (MCAo alone, remote- and penumbral-SDs) was completed in one experimental day in a randomized order, to minimize experimental bias. A grouped analysis examining the effects of experimental day and location of experimentally-induced SD on infarct size confirmed that there was no significant difference between experimental days and infarct size (two-way ANOVA, F(6, 12) = 2.808; p = 0.0604). From this analysis, the location of SD initiation maintained a statistically significant effect on infarcts 48 hours later (F (2, 12) = 29.51; p<0.0001).

These data show that SD-induction site can affect stroke outcomes, despite the fact that SDs initiated in different sites propagated similarly through infarct expansion zones. Initiation in metabolically compromised tissue, where SDs spontaneously occur clinically (Nakamura et al., 2010; Woitzik et al., 2013; von Bornstädt et al., 2015), were associated with secondary injury and lesion expansion.

### Vascular responses vary depending on location of experimental-SD initiation

Figure 4A illustrates differences in local CBF responses to SD in the infarct growth zone dependent on initiation site. Vascular responses to remote- and penumbral-SDs were monitored to examine whether there was a difference in tissue perfusion generated in response to SD. ROIs in the infarct expansion area (labeled the “growth zone”) were selected as described above and changes in local CBF perfusion were assessed in response to the last experimentally-induced SD in the cluster. Remote-SD had significantly smaller initial hypoperfusions (−9.4 ± 3.7 versus −22.0 ± 9.3 %CBF decrease, respectively; n = 6/group; p = 0.0115; Figure 4B) and subsequent small, but significant hyperperfusions compared to pre-SD baselines. Statistically, remote-SDs had significantly less hypoperfusion than penumbral SDs (8.2 ± 3.7 versus −0.39 ± 1.8 % CBF increase, respectively, n = 6/group; p = 0.0005; Figure 4C). This could suggest a mechanism where the cumulative burden of ischemia during the acute 1-hour occlusion is reduced by hyperemic responses to remote-SD initiation. Furthermore, penumbral-SD initiation caused more cumulative hypoperfusion in the growth zone during the period of dMCAo.

**Figure 4:**
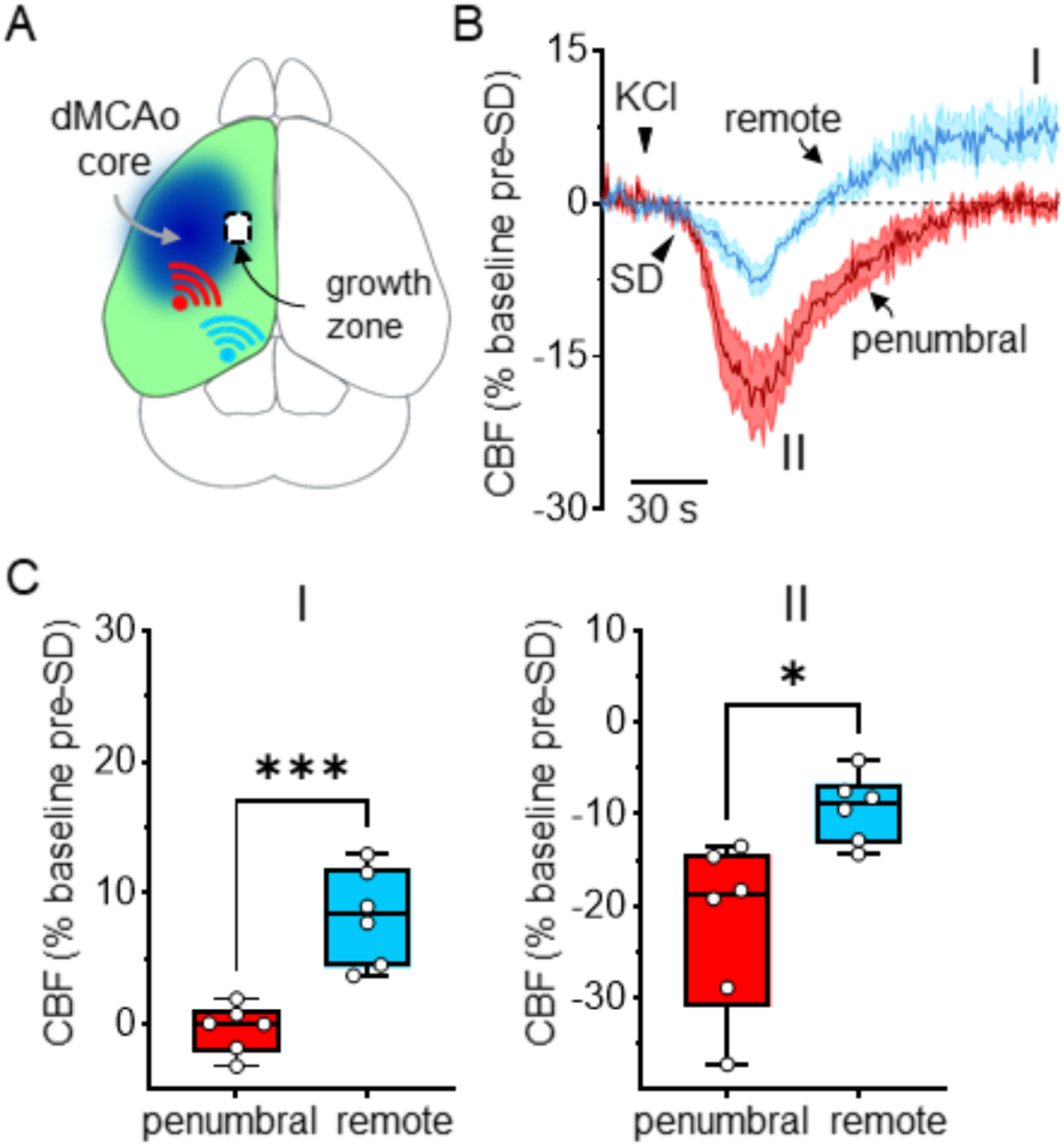
Penumbral SDs amplify ischemia. (A) Diagram showing the region termed the “growth zone” where infarct has expanded 48 hours post-occlusion. ROIs from this area were used to assess the vascular responses to SDs. (B) Traces showing CBF responses to SD from ROIs in the growth zone area in remote SD (blue trace) and penumbral SD (red trace) during the last SD in the cluster of experimentally-induced SDs. Remote SDs showed larger hyperperfusion responses following SD (I) and smaller hypoperfusion (II) (C) Summary data showing the magnitude of responses in remote SD (blue bars) and penumbral SDs (red bars). Analyses were made from the most negative deflection during SD (C_II_) and at approximately ∼2 minutes post-SD onset (C_I_) Normal distribution was evaluated by Shapiro-Wilk test (C_I_: p = 0.9766, C_II_: 0.4224). *P < 0.02, ***P < 0.0005

### Optogenetic SD stimulation show similar initiation-site mediated infarct

Finally, we repeated key studies and analyses using optogenetic stimulation of SD, rather than focal KCl stimulation. Occlusions in these experiments had similar decreases in cerebral perfusion as seen in the KCl-experiments SDs were initiated as described recently (Sugimoto et al., 2023) every 10 minutes over the 60-minute MCAo by optogenetic stimulation (3mW, 30 sec). Experimental SDs were initiated in areas as described above (Figure 6A; see Methods) and compared to MCAo alone controls. As for the KCl studies, the burden of SDs during the experimental period was from the additional SDs generated reliably from the optogenetic stimulus (average number of SDs per group: MCAo alone: 1.83 ± 0.41; remote-SD: 5.5 ± 1.9; penumbral-SD: 6.7 ± 0.81; one-way analysis of variance (ANOVA) p<0.0001; n = 6/group). Similar to results with KCl-induced SDs, with optogenetic stimulation, penumbral-SD groups had larger infarcts than both MCAo alone (16.60 ± 3.8 vs 11.71 ± 2.6 mm^3^; n = 6/group; p = 0.019; Figure 5) and remote-SD groups (16.60 ± 3.8 vs. 6.67 ± 0.48 mm^3^; n = 6/group; p < 0.0001; Figure 5). Remote-SDs were also significantly smaller than MCAo alone controls (p = 0.0159; Figure 5) suggesting that regardless of the type of SD stimulation (KCl and optogenetic) the experimental SD initiation site predicts stroke outcome.

**Figure 5:**
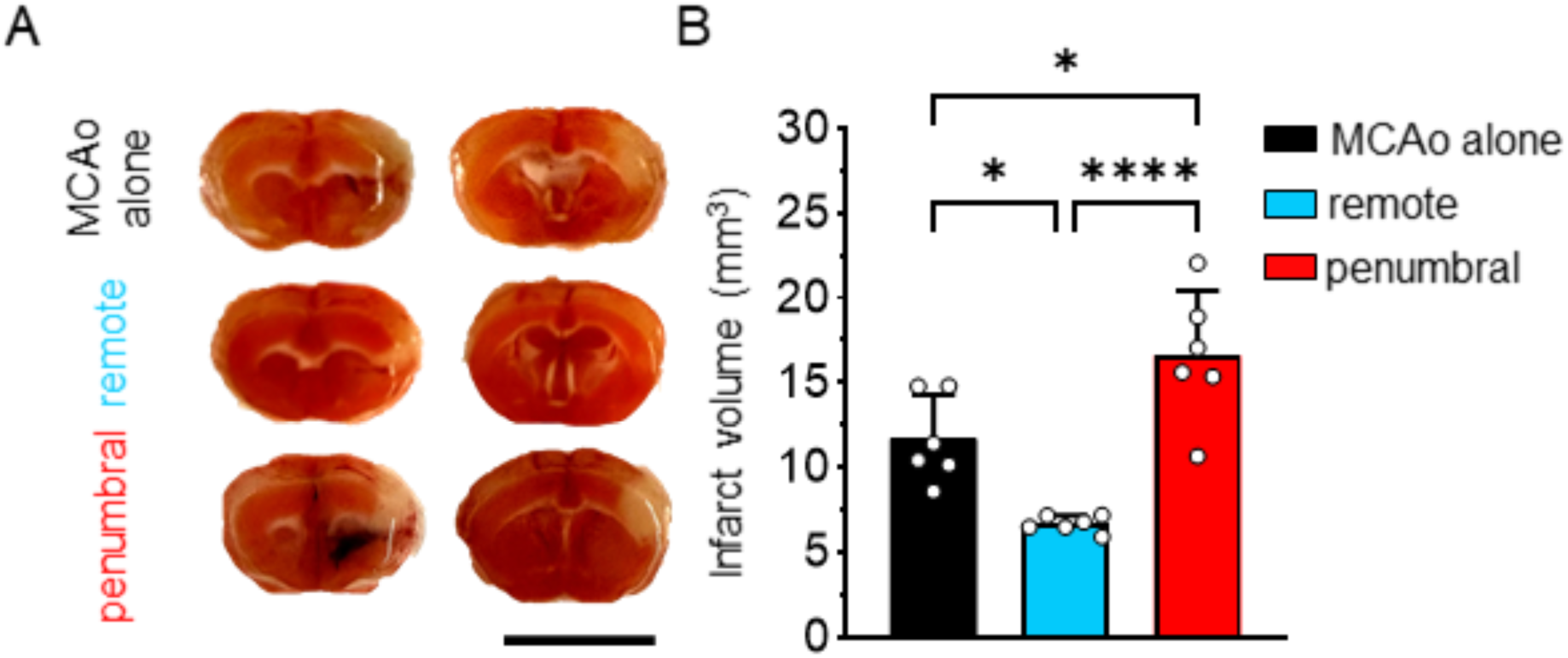
Optogenetically-initiated SDs show similar effects on infarct size. (A) Representative images of TTC stained brain sections, following (2-mm-thick). Top panel represents MCAo alone and following experimentally-generated SDs from the remote initiation site (middle row), and the penumbral initiation site (bottom row). Scale bar: 5mm. (B) Summary data of infarct sizes across the three groups: MCAo alone (black bar), remote SD (blue bar) and penumbral SD (red bar). Normal distribution was evaluated by Shapiro-Wilk test (B: p = 0.374). *P < 0.02, ****P < 0.0001

**Figure 6:**
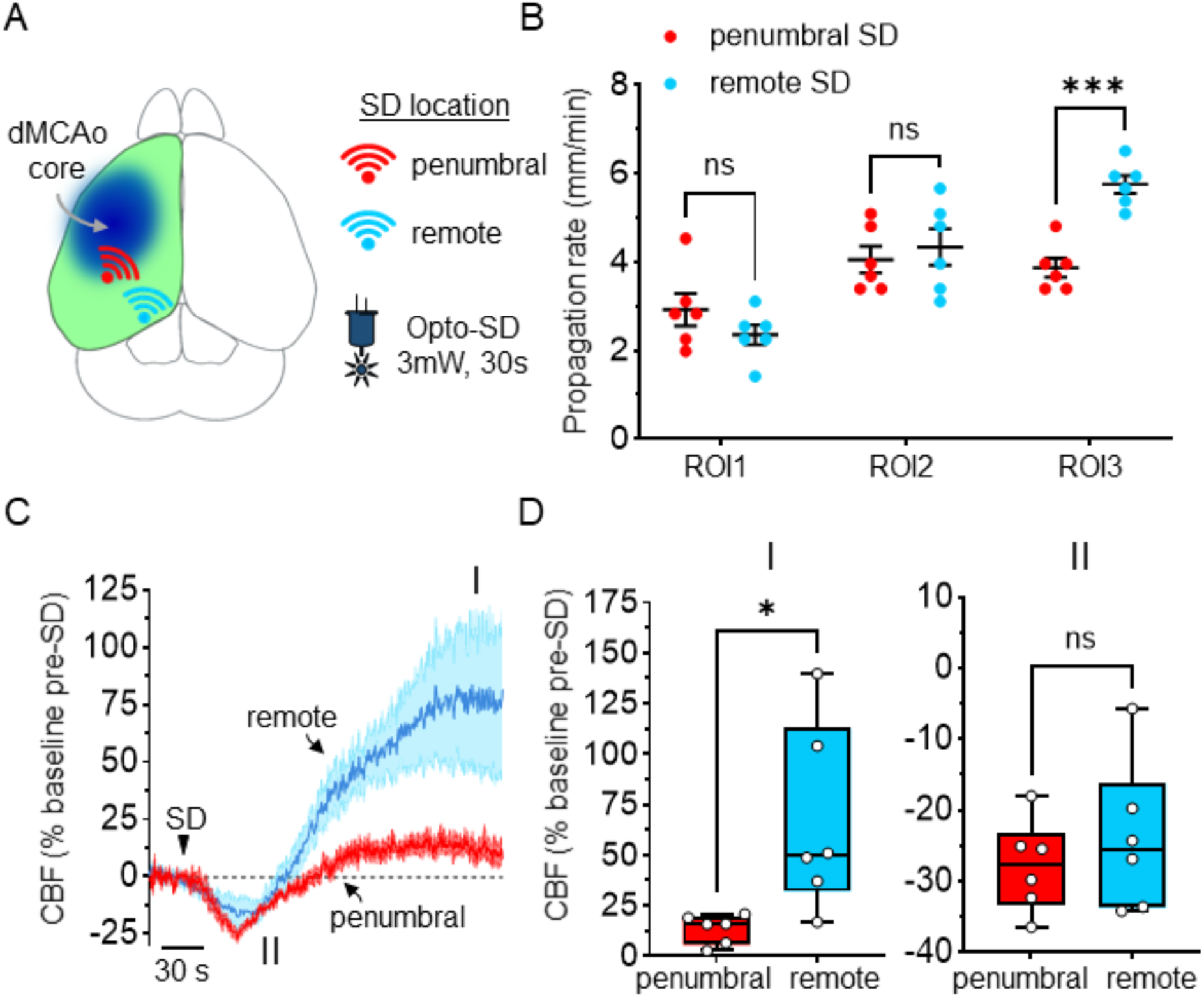
Optogenetically-initiated SDs cause similar vascular responses. (A) Diagram showing the approximate site of SD initiation in optogenetic experiments. Key showing penumbral SD group in red and remote SD group in blue. Optogenetic SDs (Opto-SD) were induced by 3 mW stimulation for 30 sec. SDs were initiated every 10 min during the first 60 min occlusion. (B) Summary data of propagation rates of Opto-SDs in both penumbral and remote SD groups showing the remote SDs only propagate faster in distal areas from the perfusion deficit (similar to Figure 3). (C) Traces of CBF perfusion during last Opto-SD in both remote SD (blue trace) and penumbral-SD (red trace) groups. (D) Summary data of the hyperperfusion (I) and hypoperfusion (II) in remote SD (blue bar) and penumbral SD (red bar). Normal distribution was evaluated by Shapiro-Wilk test (B: p = 0.419; D_I_: p = 0.277; D_II_: 0.439). *P < 0.02, ***P < 0.001

As for KCl induction, we evaluated SD propagation and vascular responses of experimentally-induced SDs. As for KCl studies, remote-SDs propagated faster than penumbral-SDs in distal regions (5.75 ± 0.496 vs. 3.87 ± 0.525 mm min^-1^; n = 6/group; p = 0.0003; Figure 6B). LSCI also imaging revealed a similar directionality of propagation where penumbral-SDs propagated first medially before invading rostral and caudal sites, (Supplemental Movie S3) whereas remote-SDs propagated uniformly in a rostral direction (Supplemental Movie S4). Regardless of initiation site, SDs propagated across the entire hemisphere. Optogenetically-initiated SDs also resulted in different vascular responses in infarct expansion zones, dependent on SD initiation site. Thus, remote-SDs had significantly larger CBF increases than penumbral-SDs (66.2 ± 46.2 vs. 13.5 ± 7.23 % CBF increase, respectively; n = 6/group; p = 0.02; Figure 6C-D). However, there was no difference in the initial hypoperfusion phase of optogenetic-induced SDs between the two groups (−24.1 ± 10.6 % CBF decrease in remote-SD vs. −27.8 ± 6.46 in penumbral-SD group; n = 6/group; p = 0.472; Figure 6C-D). Taken together with the KCl data set, these results suggest a significant contribution of SD initiation site to stroke outcomes based on a potential interplay between the length of ischemia and the amount of hyperemia. Furthermore, there was no difference in infarct development between the two SD initiation methods, with both optogenetic and KCl stimuli producing infarct enlargement when initiated at proximal sites.

## Discussion

### General findings

In this study, the side-by-side comparison of different SD initiation sites allowed demonstration of both damaging and protective effects of experimentally-induced clusters of SD in a stroke model. The findings are consistent with the conclusion that when SDs are initiated in vulnerable penumbral tissue, they are associated with increased infarct expansion (Back et al., 1996; von Bornstädt et al., 2015). A possible mechanism for injury expansion is additional cumulative oligemia in the infarct expansion zone during the period of occlusion. In contrast, SDs experimentally-initiated in more remote areas did not expand injury but were instead associated with reduced infarct sizes. This reduction was associated with improved cerebral perfusion which may decrease ischemic burden during the occlusion period. Together, these findings help resolve apparently contradictory conclusions about the roles of SDs in infarct expansion (Hartings et al., 2017; Sugimoto et al., 2023, 2024) by emphasizing the importance of sites of SD initiation sites in experimental models.

### SDs generated in peri-infarct tissue worsen acute ischemic stroke outcome

The occurrence of SDs in ischemic stroke has been strongly linked to acute secondary injury (Back et al., 1994; Busch et al., 1996; Hartings et al., 2017; Shuttleworth et al., 2020). Across species and induction methods, SDs generated in tissue lacking the metabolic supply to meet the event’s extreme demand appear to drive cortical lesion expansion (Somjen, 2001; Hartings et al., 2003; Woitzik et al., 2013; Santos et al., 2014; von Bornstädt et al., 2015; Carlson et al., 2023). Recent negative results sparked speculation that SDs are not causative in injury expansion (Sugimoto et al., 2023). Using optogenetic SD stimulation outside the perfusion deficit, they reported no effect of SDs on infarct size, suggested that neuronal stimulation used to trigger SD caused injury expansion, and concluded that the damaging role of SD in stroke should be reexamined (Sugimoto et al., 2023, 2024). The current study tested whether these negative results instead reflected the selected location of SD initiation. We therefore closely replicated their methods, using the same dMCAo model and the same timing of experimentally-evoked SDs and infarct evaluation (Sugimoto et al., 2023). Our results clearly show that stroke outcome depends on the SD-initiation site, confirming and extending evidence that SD can be injurious in stroke, while also suggesting vascular mechanisms behind these differing effects with relatively subtle experimental differences.

A first consideration is comparison of initiation sites of experimentally-induced SD with “spontaneous” SDs in injured brain. Naturally-occurring SDs are reported to initiate in peri-infarct zones and propagate around the core (Nakamura et al., 2010; Woitzik et al., 2013). In fact, sensory stimulation that results in synaptic activation in vulnerable penumbral regions (CBF <50%) presents a supply-demand mismatch that initiate SDs contributing to larger infarcts (von Bornstädt et al., 2015). These events resemble SDs initiated in proximal sites in the present study and may evoke similar vascular responses peri-infarct tissues. In the prior study, where no infarct enlargement occurred (Sugimoto et al., 2023), SDs were initiated in peri-infarct areas, but without reporting of the precise location with respect to perfusion deficit of dMCAo. This raises the possibility that the SDs were initiated in intermediate regions between the sites defined in our study. Such a location would be expected to produce effects between the damaging and beneficial outcomes observed here, consistent with no significant change in infarct size reported in that study (Sugimoto et al., 2023).

A second issue concerns the stimulus used to initiate SD. It was suggested (Sugimoto et al., 2023) that damaging effects of SD in prior reports reflected noxious effects of KCl stimuli rather than any deleterious effects of SD itself. The lack of infarct growth was attributed to optogenetic stimulation being less invasive, irrespective of SD generation (Sugimoto et al, 2023). The present results suggest that this is unlikely to be the case. Both damaging and protective SD effects were observed with KCl stimulation, differing only by SD initiation site. An unlikely result if KCl applications are themselves toxic in this model, and consistent with reports lacking damaging effects of KCl-induced SDs (Nedergaard and Hansen, 1988). Moreover, SDs generated with the less invasive optogenetic method also demonstrated both damaging and beneficial effects, depending on initiation site. Thus, optogenetic activation of SD cannot be considered intrinsically less damaging in this model. Results with both methods support the conclusion that location of SD initiation, not type of stimulus, determines the burden of SD on stroke outcomes.

Optogenetic activation of peri-infarct tissue (without generation of SD) has been linked to infarct enlargement (Sugimoto et al., 2024). These observations complement literature documenting varied effects of functional activation on infarct size and tissue recovery. Across experimental designs, synaptic stimulation in peri-infarct zones (without intentional activation of SD) improved cerebral blood flow to the ischemic area and was neuroprotective (Davis et al., 2011; Lay et al., 2011). However, this neuroprotection was restricted to the first 2 hours post-MCA occlusion, where stimulation beyond 3 hours worsened outcomes (Lay and Frostig, 2014). Together, these studies suggest a context-dependent effect of neural stimulation in ischemic stroke that exists below the SD threshold.

### Vascular response is dependent on the location of SD induction

It is unlikely that the properties of SDs themselves are fundamentally different when initiated in proximal or remote locations. While we observed differences in propagation directionality, likely due to the territory available for SD invasion, the rate of propagation was only significantly different between penumbral- and remote-SDs in distal, less compromised areas. Rates in peri-infarct tissues and infarct growth areas was not significantly different. Similar propagation patterns between the KCl-SD and optogenetic-SD groups further suggest that the SD wave was not different across initiation sites. Our data instead suggest variation in vascular responses in infarct expansion zones, determined by the location of SD initiation and SD wave directionality. Increases in the propagation rate in distal regions could be exaggerated by collateral blood flow from adjacent vascular territories. The growth zone showed no such effect, suggesting that spreading hyperemia as a surrogate for SD propagation could be confounded by the vascular anatomy and tissue metabolic status.

A similar directionality was observed in a study examining vascular responses to spontaneous SD in ischemic stroke (Binder et al., 2022). They identified two consequences of spontaneous SDs. SDs that emerged from the penumbra and propagated outward produced larger infarcts, and SDs that invaded the entire hemisphere had significantly smaller infarcts, than SD naïve mice (Binder et al., 2022). Our results showed similar propagation of experimental-SDs but identified that the stroke outcome depends on the site of SD initiation, not on the extent of territory it invades.

In our study, clusters of remote-SDs enhanced cerebral perfusion during and after SD. Specifically, within the infarct growth zone, initial oligemic hypoperfusion was small, and subsequent hyperperfusion was not dissimilar from vascular responses to SD in healthy cortex (Ayata and Lauritzen, 2015). This suggests SD-mediated (likely retrograde) vascular flow to the ischemic territory. Collateral circulation through pial arterial networks can improve tissue outcomes from focal stroke (Riva et al., 2012; Cuccione et al., 2016). Remote-SDs were initiated within the territory of the anterior cerebral artery (ACA), and it should be noted that the growth zone was near the ACA-MCA watershed zones. Thus, remote-SDs could promote collateral flow and metabolic support to peri-infarct zones normally primed for injury expansion. While interesting, this method of experimental SD-induction is far removed from spontaneous SDs and thus any neuroprotective effects of SD are not expected during the normal clinical course of stroke patients in the ICU. These data do raise the question of whether intentional SD-stimulation in remote cortex, using an approach such as transcranial magnetic stimulation, could replicate these neuroprotective effects in stroke patients.

Alternatively, penumbral-SD clusters showed a more profound oligemia in the same growth zone. While hypoperfusion eventually normalized, it never transitioned to significant hyperperfusion. Capillary dysfunction causes neurovascular uncoupling in regions surrounding the ischemic core and has been linked to spontaneous SDs (Staehr et al., 2023). Furthermore, consecutive spontaneous SDs after photothrombotic stroke showed progressive worsening of tissue vascular responses (Zhao et al., 2021). Our data reinforce the idea that penumbral-SDs cause injury expansion (von Bornstädt et al., 2015; Reinhart et al., 2023), but highlight underrecognized vascular responses which predict stroke outcome. Our findings propose prolonged ischemic burden in peri-infarct tissue due to SD-mediated vascular dysfunction. Thus, future studies should rigorously select SD stimulations that model spontaneous penumbral events to avoid confounding vascular responses.

### Conclusion

This study helps to address the distinction between correlation and causation of SD in the progression of stroke injury. Side-by-side comparison of different SD stimuli and initiation locations reveals that the site of SD initiation (rather than stimulus type) can result in very different stroke outcomes. The findings strengthen conclusions that SDs can be causative in the progression of injury and support efforts to clinically target SDs that arise from vulnerable brain regions in stroke and trauma patients. Furthermore, the results emphasize the importance of SD initiation sites in experimental stroke and brain injury studies.

## CRediT Author Contributions

**Michael C. Bennett:** Conceptualization, Data curation, Formal analysis, Funding acquisition, Investigation, Visualization, Writing – original draft. **Russell A. Morton:** Conceptualization, Writing – review and editing. **Andrew P. Carlson:** Conceptualization, Writing – review and editing. **C. William Shuttleworth:** Conceptualization, Funding acquisition, Methodology, Project administration, Supervision, Writing – review & editing.

## Acknowledgements

We thank the UNM Preclinical Core of the Center from Brain Recovery and Repair for technical support. Supported by NIH grants NS129351, NS106901, GM109089 & GM159568. Schematics in Figs 1B, 2A, 4A, and 6A were created with Biorender.

## Supplemental Video Legends

**Video S1: SD initiated in the penumbral area by KCl**

Representative pseudo-colored LSCI video of a propagating SD initiated in the penumbral region (CBF decrease >50% of baseline) by focal KCl application (see Methods). Video is at 25x speed for visualization of the propagating SD. Scale: 2 mm

**Video S2: SD initiated in the remote area by KCl**

Representative pseudo-colored LSCI video of a propagating SD initiated in the remote region (CBF decrease <30% of baseline) by focal KCl application (see Methods). Video is at 25x speed for visualization of the propagating SD. Scale: 2 mm

**Video S3: SD initiated in the penumbral area by optogenetic stimulation**

Representative pseudo-colored LSCI video of a propagating SD initiated in the penumbral region (CBF decrease >50% of baseline) by optogenetic stimulation (see Methods). Video is at 25x speed for visualization of the propagating SD. Scale: 2 mm

**Video S4: SD initiated in the remote area by optogenetic stimulation**

Representative pseudo-colored LSCI video of a propagating SD initiated in the remote region (CBF decrease <30% of baseline) by optogenetic stimulation (see Methods). Video is at 25x speed for visualization of the propagating SD. Scale: 2 mm

